# Human gut bacteria tailor extracellular vesicle cargo for the breakdown of diet- and host-derived glycans

**DOI:** 10.1101/2023.04.03.535451

**Authors:** Mariana G. Sartorio, Evan J. Pardue, Nichollas E. Scott, Mario F. Feldman

**Affiliations:** Department of Molecular Microbiology, Washington University School of Medicine, Saint Louis, MO, 63110, USA; Department of Microbiology and Immunology, The Peter Doherty Institute for Infection and Immunity, University of Melbourne, Parkville, Victoria, Australia

## Abstract

Extracellular vesicles (EV) are produced in all three domains of life, and their biogenesis have common ancient origins in eukaryotes and archaea. Although bacterial vesicles were discovered several decades ago and multiple roles have been attributed to them, no mechanism has been established for vesicles biogenesis in bacteria. For this reason, there is a significant level of skepticism about the biological relevance of bacterial vesicles. In *Bacteroides thetaiotaomicron* (*Bt*), a prominent member of the human intestinal microbiota, outer membrane vesicles (OMVs) have been proposed to play key physiological roles. By employing outer membrane- and OMV-specific markers fused to fluorescent proteins we visualized OMV biogenesis in live-cells. We performed comparative proteomic analyses to demonstrate that *Bt* actively tailors its vesicle cargo to optimize the breakdown of diet- and host-derived complex glycans. Surprisingly, our data suggests that OMV are not employed for mucin degradation. We also show that, in *Bt*, a negatively-charged N-terminal motif acts as a signal for protein sorting into OMVs irrespective of the nutrient availability. We conclude that OMVs are the result of an exquisitely orchestrated mechanism. This work lays the foundation for further investigations into the physiological relevance of OMVs and their roles in gut homeostasis. Furthermore, our work constitutes a roadmap to guide EV biogenesis research in other bacteria.

## Introduction

The human gastrointestinal tract harbors one of the densest microbial communities known in nature. A healthy adult human gut microbiota is primarily colonized by two bacterial phyla: Firmicutes and Bacteroidota^1, 2^. *Bacteroides* spp. are one of the most abundant genera, representing nearly one-third of the total human gut microbiota^3^. It has been shown that *Bacteroides* spp. promote gut homeostasis by shaping host immunity^4–6^, and preventing pathogen colonization^7, 8^. Moreover, *Bacteroides* is well-known as glycan-degrading specialist that can metabolize a wide array of polysaccharides, derived from host glycans and dietary fibers^9–12^. This metabolic flexibility resides in a series of gene clusters termed <u>P</u>olysaccharide <u>U</u>tilization <u>L</u>oci (PULs), co-localized and co-regulated loci in response to specific glycans that enable the degradation and uptake of specific carbon sources^13^. Recent studies show that *Bacteroides* spp. produce outer membrane vesicles (OMV) that participate in glycan utilization and the modulation of host immune responses in the gut^6, 14, 15^.

In Gram-negative bacteria, OMV are generated by the blebbing of the outer membrane (OM). OMVs are composed mainly of phospholipids, lipopolysaccharide or lipooligosaccharide, as well as periplasmic and membrane proteins^16^. Numerous bacterial processes have been proposed to be mediated by OMVs, including envelope stress responses, delivery of virulence factors, modulation of host immune responses, and digestion of extracellular nutrients^16^. Despite being discovered more than 50 years ago^17, 18^, the field has struggled to progress due to three crucial flaws: (1) there is no method to visually distinguish between genuine OMVs from lysis byproducts that occur during normal cell growth, (2) no reliable ways to quantify OMV production under physiologically relevant conditions are available, and (3) no general mechanism for OMV biogenesis has been established. For these reasons, the microbiology community remains partially skeptical about the biological significance of the OMVs^19^.

Previous mass spectrometry (MS) analyses have demonstrated that, in *Bacteroides thetaiotaomicron* (*Bt*) and *Bacteroides fragilis* (*Bf*), a subset of proteins is selectively targeted to OMVs, while others are retained in the OM and not directed to vesicles^20, 21^. Furthermore, many OMV-localized proteins were shown to be lipoproteins that share common functionalities (i.e. glycosylhydrolases and proteases), and contain a negatively charged “<u>L</u>ipoprotein <u>E</u>xport <u>S</u>ignal (LES)” motif that was shown to be necessary for surface localization and possibly to efficiently targeting them to OMVs^21, 22^. In addition, it has been shown that OMVs from *Bacteroides spp.* retain glycolytic activity and can be used to supplement the growth of bacteria on carbon sources they normally could not utilize ^23, 24^. To the best of our knowledge, these features are exclusively found in *Bacteroides*-derived vesicles.

In this work, we employed live-cell fluorescence microscopy to visualize for the first time OMV biogenesis, and performed comparative proteomic analyses to demonstrate that *Bt* actively alters the content of their OMVs to gain a fitness advantage in different growth conditions. Our results show that the production of OMVs in *Bt* is the result of a highly regulated process.

## Results

### Vesicle cargo proteins localize at defined foci on the bacterial membrane

Due to the unique properties, we hypothesized that *Bacteroides* OMV production is the result of an orchestrated cellular process. To test this hypothesis, we exploited the differential sorting of proteins into OMVs to visualize their biogenesis by live fluorescence microscopy. First, we identified proteins that were either enriched in OMVs or retained in the OM, and fused them to sfGFP or mCherry. As OMV markers we chose three different lipoproteins previously reported to be preferentially packed in vesicles: BACOVA_04502 (inulinase) from *Bacteroides ovatus* (*Bo*)^20, 23, 25^, BF_1581 from *Bf*^20^ and BT_3698 (SusG) from *Bt*^21^. As OM markers we chose the integral OM protein BT_0418 (OmpF) and the lipoprotein BT_2844^21^. The chimeric versions of all candidates were cloned under constitutive promoters into the pNBU2 integrative vector in *Bt* (see Table S1 and materials and methods). Western blot analysis of subcellular fractions demonstrated that the fusion of fluorescent proteins did not alter the partitioning between OM and OMVs of each construct (fig. 1A, fig. S1).

**Figure 1.**
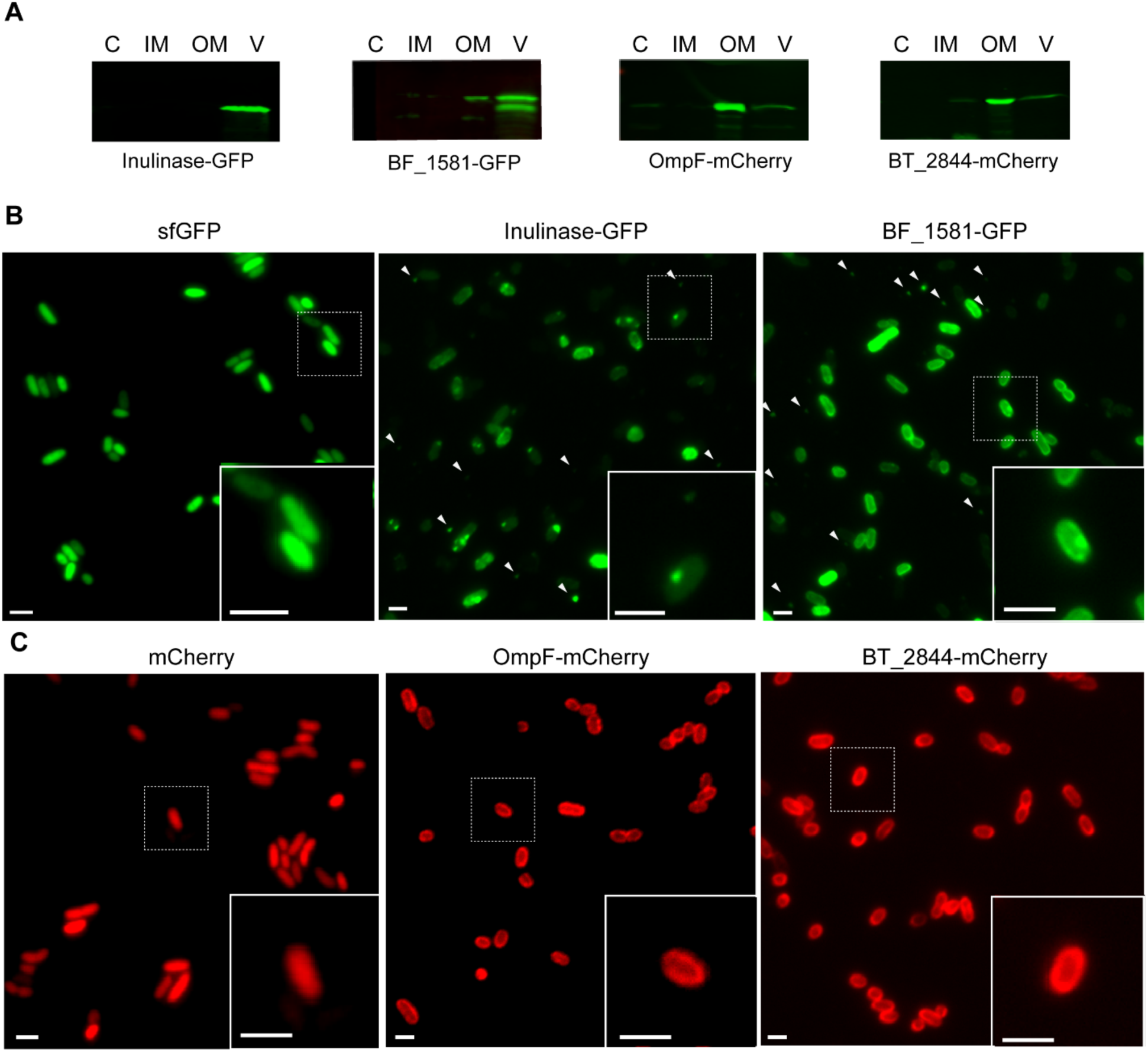
OMV and OM chimeric markers show a differential distribution in *Bt*. **A.** Western blot of 10 μg of protein from soluble fraction (S), inner membrane (IM), outer membrane (OM) and OMV (V) fractions of *Bt* expressing Inulinase-GFP, BF_1581-GFP, OmpF-mCherry or BT_2844-mCherry. Anti-His and anti-mCherry antibodies were employed to identify GFP and mCherry chimeric markers, respectively. **B.** Representative widefield fluorescence microscopy images of OMV chimeric markers Inulinase-GFP and BF_1581-GFP. White arrows indicate the presence of OMVs. **C.** Representative widefield fluorescence microscopy images of OM chimeric markers OmpF-mCherry and BT_2844-mCherry. Cytosolic expression of sfGFP and mCherry are used as reference (left panels). Scale bar: 2 μm.

*Bt* is an anaerobe, however, both mCherry and sfGFP require oxygen for fluorophore maturation. It has been proposed that this limitation can be overcome by exposing bacteria to nanaerobic concentrations of oxygen prior to their visualization by fluorescence microscopy^25^. Although in these conditions we were able to visualize GFP fluorescence, our attempts to visualize chimeric proteins containing mCherry were unsuccessful. However, proper fluorophore maturation and fluorescence was achieved after exposure to aerobic conditions (see materials and methods). Although atmospheric concentrations of oxygen are not physiological for *Bacteroides spp*., neither cell viability nor the distribution of our chimeric markers was affected after 4 hours of exposure to air (fig. S2). In all cases, fluorescent chimeric proteins were detected at the cell surface. We frequently observed the OMVs markers localized at defined foci (fig. 1B, inset). This phenotype was not always present, and the number of foci was variable. On the contrary, OM markers invariably exhibited a more homogeneous distribution along the cell borders (fig. 1C, inset). Interestingly, round-shaped extracellular structures of a size compatible with vesicles were observed in *Bt* expressing OMV chimeric markers inulinase, BT_1581 or SusG fused to sfGFP (fig. 1B, fig. S3). These structures were not detected in the cells expressing the OM chimeric markers OmpF and BT_2844 fused to mCherry (fig 1C). These results suggest that chimeric fluorescence proteins can be employed to visualize OMVs and we hypothesize that the punctate distribution observed for the OMV markers is related to the recruitment of vesicle-targeted proteins to specific foci prior to vesicle formation.

### Visualizing OMV formation

We subsequently analyzed whether the co-expression of an OMV marker along with an OM marker enables the distinction between OMVs and cells or lysis byproducts by fluorescence microscopy. We reasoned that lysis byproducts would be generated by the non-specific breakage of cellular membranes, and, due to this, should contain both, OM and OMV markers. On the contrary, genuine OMVs are not expected to contain OM markers and, instead, should only be detected via the proper OMV markers. Western blot analysis confirmed that co-expression of the different chimeric proteins did not affect their original OMV versus OM distribution (fig. S4). *Bt* cells co-expressing Inulinase-GFP and OmpF-mCherry were detected in both, green and red fluorescence channels. Remarkably, extracellular structures were exclusively detected in the green channel (fig. 2A, white arrows). Furthermore, when we simultaneously expressed two different OMV markers, Inulinase-GFP and BF_1581-mCherry, both co-localized in the extracellular structures (fig. 2B, white arrows). Thus, together our results dismiss the notion that OMVs result from bacterial lysis.

**Figure 2.**
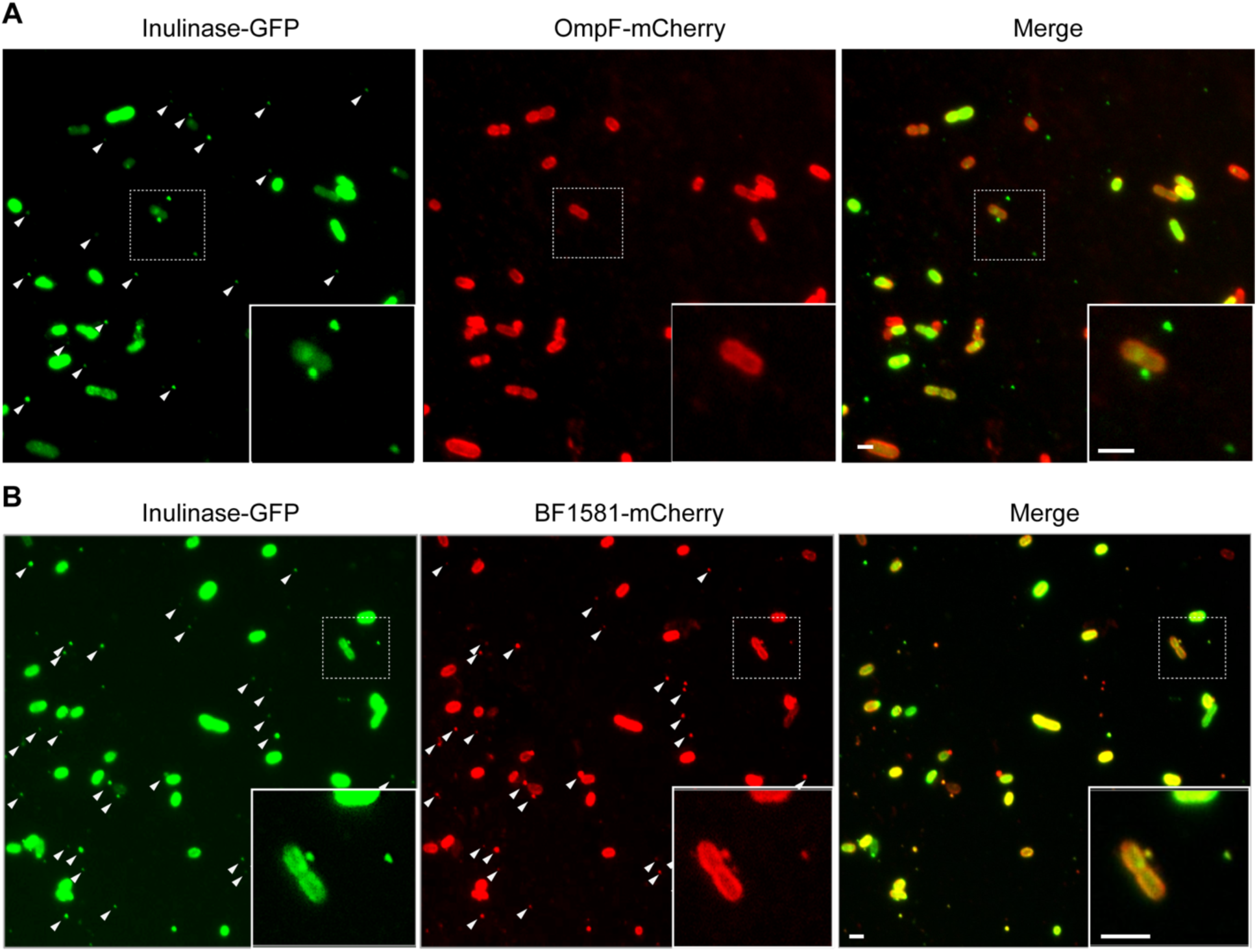
Co-expression of OMV and OM markers do not show co-localization in vesicles. Representative widefield fluorescent microscopy images of *Bt* co-expressing **A.** Inulinase-GFP and OmpF-mCherry, or **B.** Inulinase-GFP and Bf_1581-mCherry. Scale bar: 2 μm.

To capture OMV biogenesis in live cells, we utilized time lapse widefield fluorescent microscopy of *Bt* co-expressing Inulinase-GFP and OmpF-mCherry. Here, we observed that the OM marker invariably remained in the cell, while the OMV markers were found to be released from the cell (fig. 3, white arrows; supplementary movie S1). It was also observed that some OMVs were not released by the bacteria (fig. 3, blue arrows, supplementary video S1), evidencing that OMV cargo proteins have dynamic distributions over time. These results show, for the first time, vesicle biogenesis dynamics in live bacteria. Altogether, the observations support the existence of OMV protein sorting mechanisms as a part of an orchestrated process in *Bacteroides* spp.

**Figure 3.**
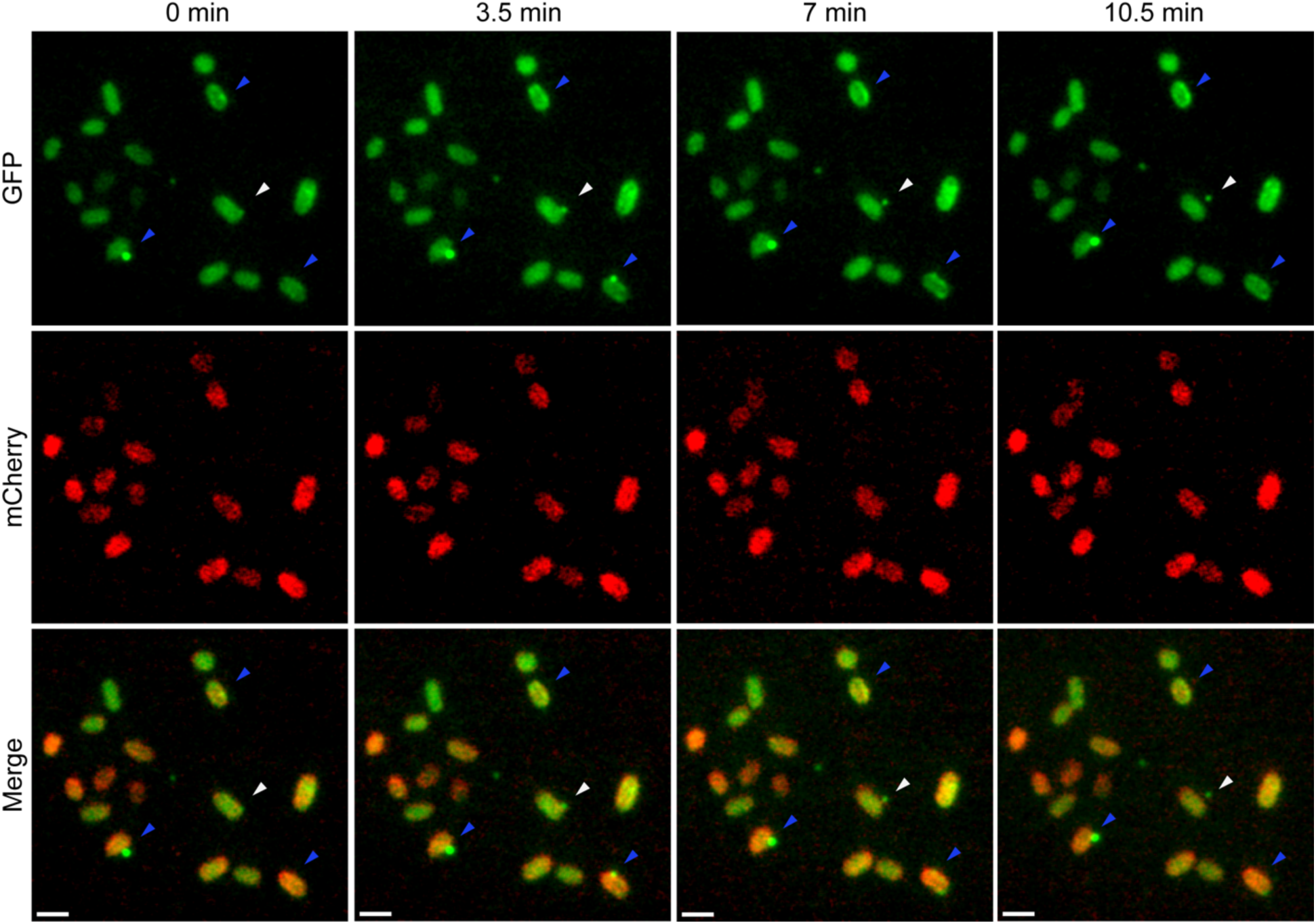
Time-lapse of OMV formation. *Bt* co-expressing Inulinase-GFP and OmpF-mCherry was grown overnight in anaerobic conditions in liquid minimal media supplemented with glucose. Cultures were then incubated at 37 °C in the presence of oxygen for 3 hours for fluorophore maturation. Cultures were diluted in pre-warmed minimal media supplemented with glucose and loaded onto 1% agarose pads for image acquisition. Images were acquired in a 37 °C pre-warmed microscope every 3.5 min. Scale bar: 2 μm. (See supplementary video S1).

### Proteomic analysis of membrane and OMV fractions

*Bacteroides* is highly specialized in the degradation of host and diet-derived glycans ^9–12^. We have previously shown that OMVs are equipped with numerous glycosyl hydrolases and other degradative enzymes required for the digestion of nutrients^20, 21^. We hypothesized that *Bt* can modulate the protein cargo of their OMVs to effectively utilize diverse array of glycans. To test this, we cultured *Bt* in minimal media supplemented with different polysaccharides or glycosaminoglycans: levan, starch, mannan, hyaluronan, heparin or mucin; glucose was employed as a reference condition. SDS-PAGE of OM and OMV fractions from *Bt* grown under the different conditions revealed significant variations in protein profiling (fig. 4). Surprisingly, the amount of protein detected in OMV fractions from cells culture in mucin and glucose were significantly lower than in the other conditions.

**Figure 4.**
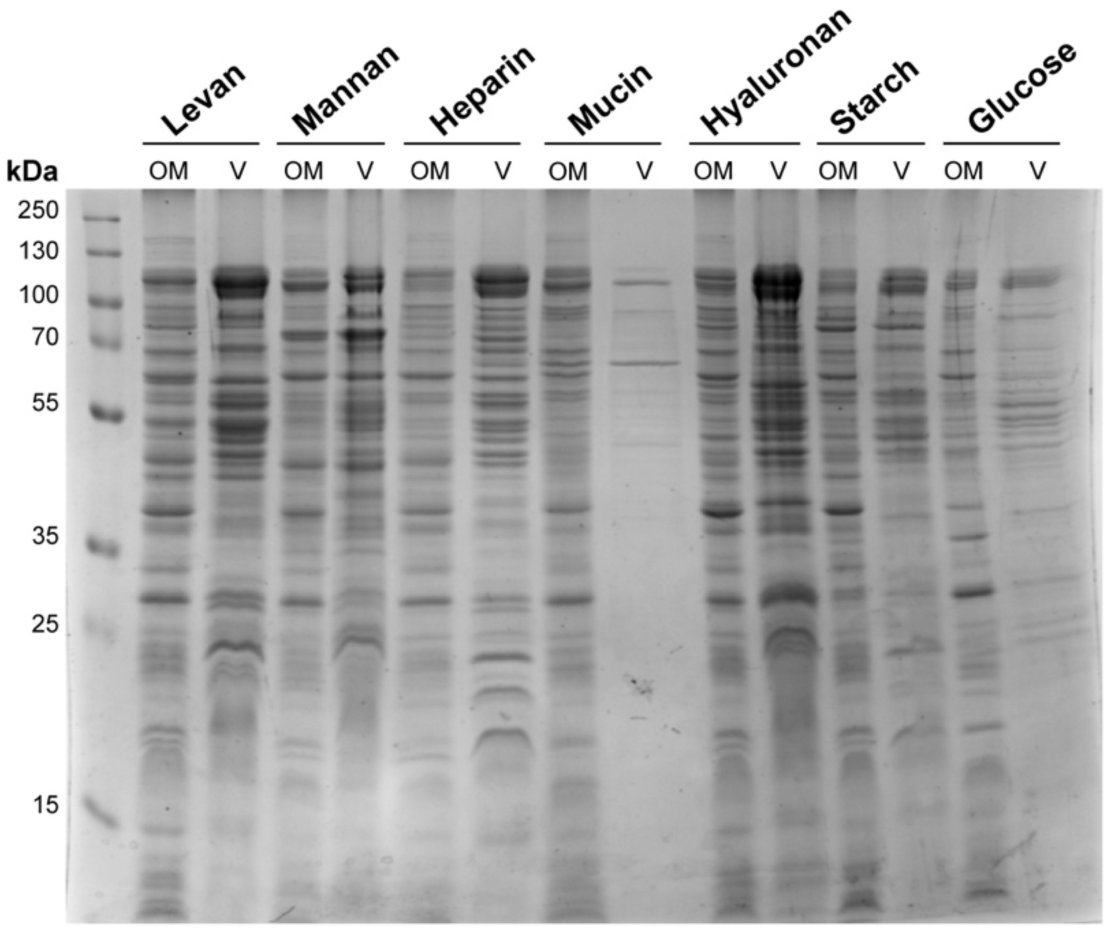
*Bt* modulates OMV cargo in different nutrient conditions. Coomassie blue staining after SDS-PAGE of *Bt* OM and OMV (V) fractions obtained after growth in minimal media supplemented with levan, mannan, heparin, mucin, hyaluronan, starch or glucose. Fractions were normalized by OD (600 nm).

To further analyze their composition, we performed comparative proteomic analyses of OMVs and OMs from *Bt* cultured in all these nutrients. Principal component analysis (PCA) of OMVs and OM proteomics showed that the biological replicates cluster together (fig. S5). This clustering was also observed by Pearson correlation analysis, with correlation coefficients of above 0.85 in most of the cases (fig. S6), revealing highly similar biological replicates. When comparing OMV proteomic datasets, we observed higher correlations between samples obtained after growth in levan and starch, as well as between heparin and hyaluronan, indicating that the proteome is more similar between these groups (fig. S6). Instead, OMVs obtained under growth in mucin were the most divergent ones, displaying the lowest correlation coefficients when compared to all the conditions (fig. S6). This is also reflected in the PCA, where the mucin proteome clusters separately (fig. S5). Remarkably, this divergent behavior was not observed in the OM comparative proteomic analysis, where OM proteome from *Bt* grown in mucin is found to be more similar to the OM proteome from *Bt* grown in hyaluronan and heparin (fig. S5 and S6). This suggests that the changes in the content of the OMVs do not necessarily correspond with the changes in the OM, likely as a result of a differential protein sorting into vesicles depending on the glycan available.

### Lipoproteins containing the negatively-charged LES signal are specifically packed in OMVs of bacteria grown in all conditions

We previously observed that, when *Bt* was cultured in rich media, most OMVs-enriched lipoproteins contain the N-terminal motif S(D/E)_3_^21^. Moreover, mutations in this motif in the lipoprotein SusG abrogates its sorting into OMVs^21^. A similar motif, K-(D/E)_2_ or Q-A-(D/E)_2_, named the LES motif, was proposed by Lauber and colleagues to be required for lipoproteins surface exposure in the OM in the oral pathogen *Capnocytophaga canimorsus,* another member of the Bacteroidota phylum^22^. Through our comparative proteomics experiments we tested if the LES-based partition occurred in presence of seven different carbohydrate nutrients. Volcano plots presented in figure 5A indicate that, in all conditions, a great majority of the proteins preferentially sorted into OMVs contain the LES motif (labelled in red in figure 5A, fig. 5B, tables S2-S8). On the contrary, integral components of the membrane were mostly retained in the OM fraction (labelled in blue). Additionally, we found that lipoproteins lacking the LES motif (labelled in yellow in fig. 5A) are mostly retained in the OM (fig. 5A), irrespective of the glycan present in the media. These experiments strongly support a model in which the LES motif acts as a conserved signal for lipoprotein sorting into OMVs.

**Figure 5.**
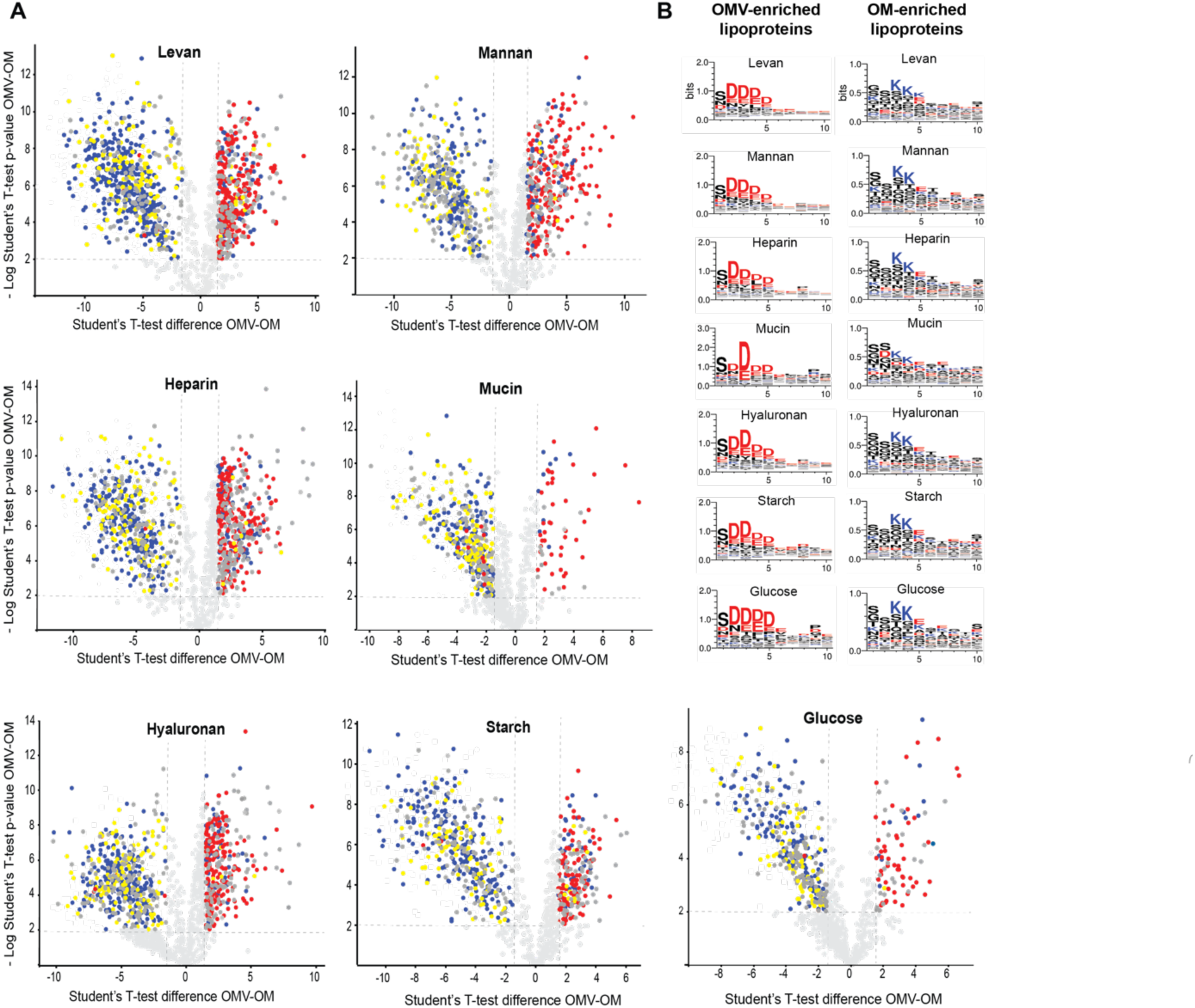
OMV-enriched lipoproteins expressed in different culture conditions harbor a conserved N-terminal LES motif**. A.** Volcano plot representations of OM and OMV-enriched proteins. Integral membrane proteins are represented in blue, lipoproteins with LES motifs are indicated in red, lipoproteins lacking the LES motif are depicted in yellow, and soluble proteins are indicated in dark gray. **B.** Lipoprotein amino acid residues next to the cysteine required for acylation were aligned. Left panels shows prevalent amino acid residues in OMV-enriched lipoproteins (cut-off: OMV/OM fold change >2) and right panels shows prevalent amino acid residues in OM-enriched lipoproteins (cut-off: OMV/OM fold change <-2). Negatively-charged amino acid residues are colored in red, and positively-charged in blue. The consensus sequences were generated using WebLogo (https://weblogo.berkeley.edu/logo.cgi).

### OMV cargo is tailored for the digestion of specific polysaccharides

Comparative proteomic analysis between OMVs from *Bt* grown in the different conditions determined that specific subsets of proteins are preferentially sorted into vesicles depending on the glycan present in the media (fig. 6, red hits; table S9). Proteins that are underrepresented in the OMVs for each growth condition are also illustrated (fig. 6, blue hits). Notably, although the OMV-enriched proteins vary for each glycan, most of these proteins are functionally and, in many cases, genetically related. Many of the proteins are glycosyl hydrolases encoded within PULs, which contain proteins involved in the binding, uptake and degradation of specific polysaccharides that are co-regulated in response to specific nutrients to enable glycan utilization. For example, proteins belonging to the heparin utilization PUL 85, such as BT_4652, BT_4675 and BT_4662 were particularly enriched in OMVs produced by cells cultured in presence of heparin, but not in the presence of the other glycans. Similarly, proteins encoded in PUL 57, proposed to be involved in hyaluronan and chondroitin sulphate degradation, were enriched in OMVs produced by bacteria grown in hyaluronan (table S9). Most of the induced PULs identified in our proteomic analysis correlate with findings from previous transcriptomic studies performed with *Bt in vivo* and *in vitro*^9^. Additionally, we identified that components of unassigned PULs were specifically induced and enriched in OMVs in response to specific nutrients. For example, protein components of PULs 41 and 42 are specially induced and targeted to OMVs in presence of heparin (Table S9).

**Figure 6.**
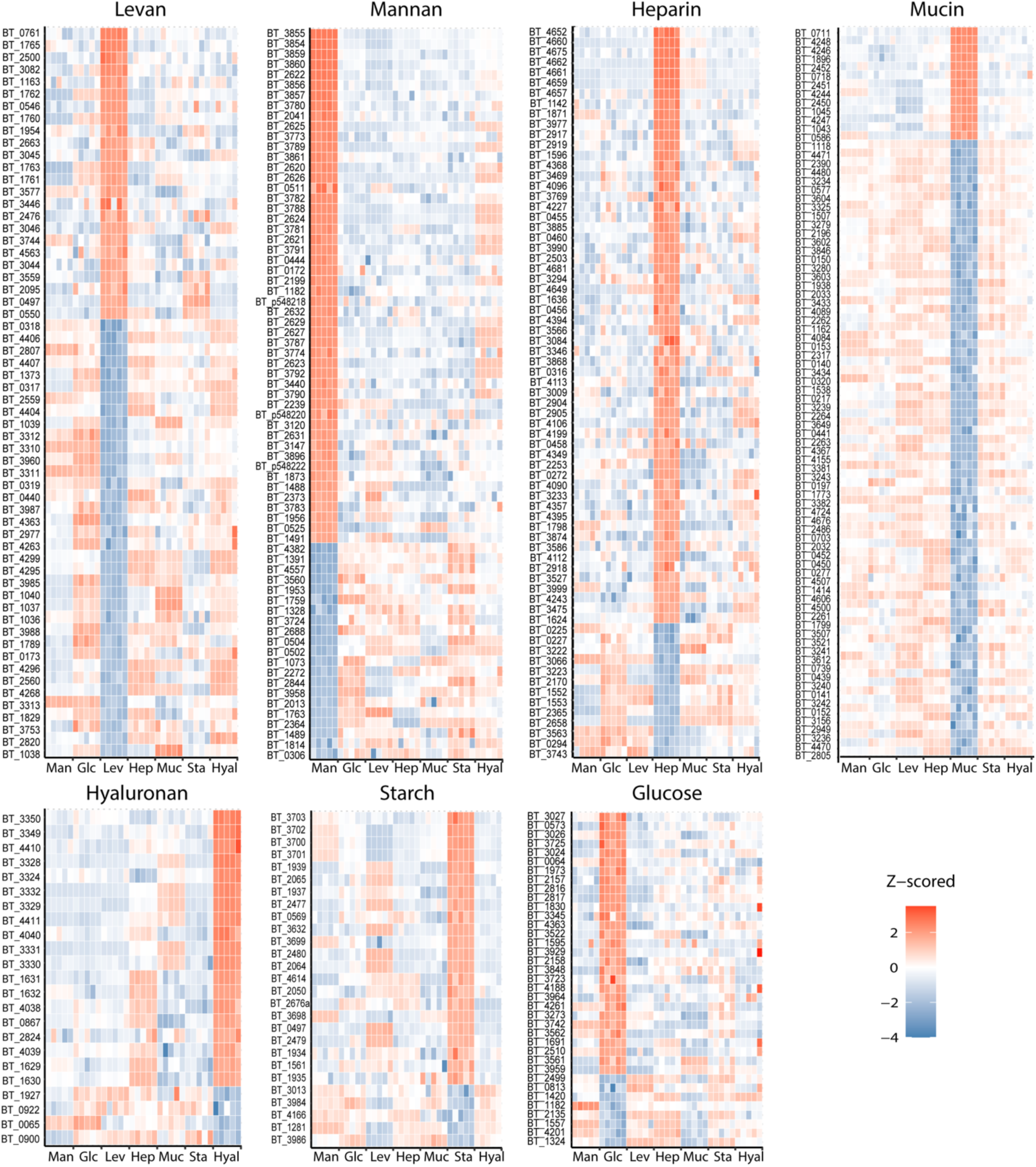
OMV cargo is tailored for the digestion of specific polysaccharides. Heat maps of protein levels (Z-score) showing most enriched (red) and excluded (blue) proteins found in OMVs from *Bt* grown in minimal media supplemented with levan (Lev), mannan (Man), heparin (Hep), mucin (Muc), hyaluronan (Hyal) or glucose (Glc) as carbon sources. Five biological replicates were performed per growth condition.

Surprisingly, despite mucin being the most complex carbon source employed in these experiments, mucin-derived OMVs displayed a reduced number of proteins targeted to vesicles (fig. 6). However, the comparative OM proteome dataset indicates that the PULs previously reported to be required for mucin degradation are indeed expressed (table S10). In addition, OM fractions from bacteria grown in mucin displayed a higher number of LES-containing lipoproteins in comparison to all other growth conditions (Table S5). This suggests that proteins that would usually be targeted to the OMVs are retained in the presence of mucin. Figure 7 illustrates the behavior of PUL components in response to different nutrients. PUL-derived proteins were enriched in the OMV fraction for all the conditions except for mucin (green circles). In fact, in presence of mucin, PUL-derived proteins were either retained in the OM (orange circles) or equally distributed in both fractions (gray circles). The complete PUL dataset analysis is presented in figure S7.

**Figure 7:**
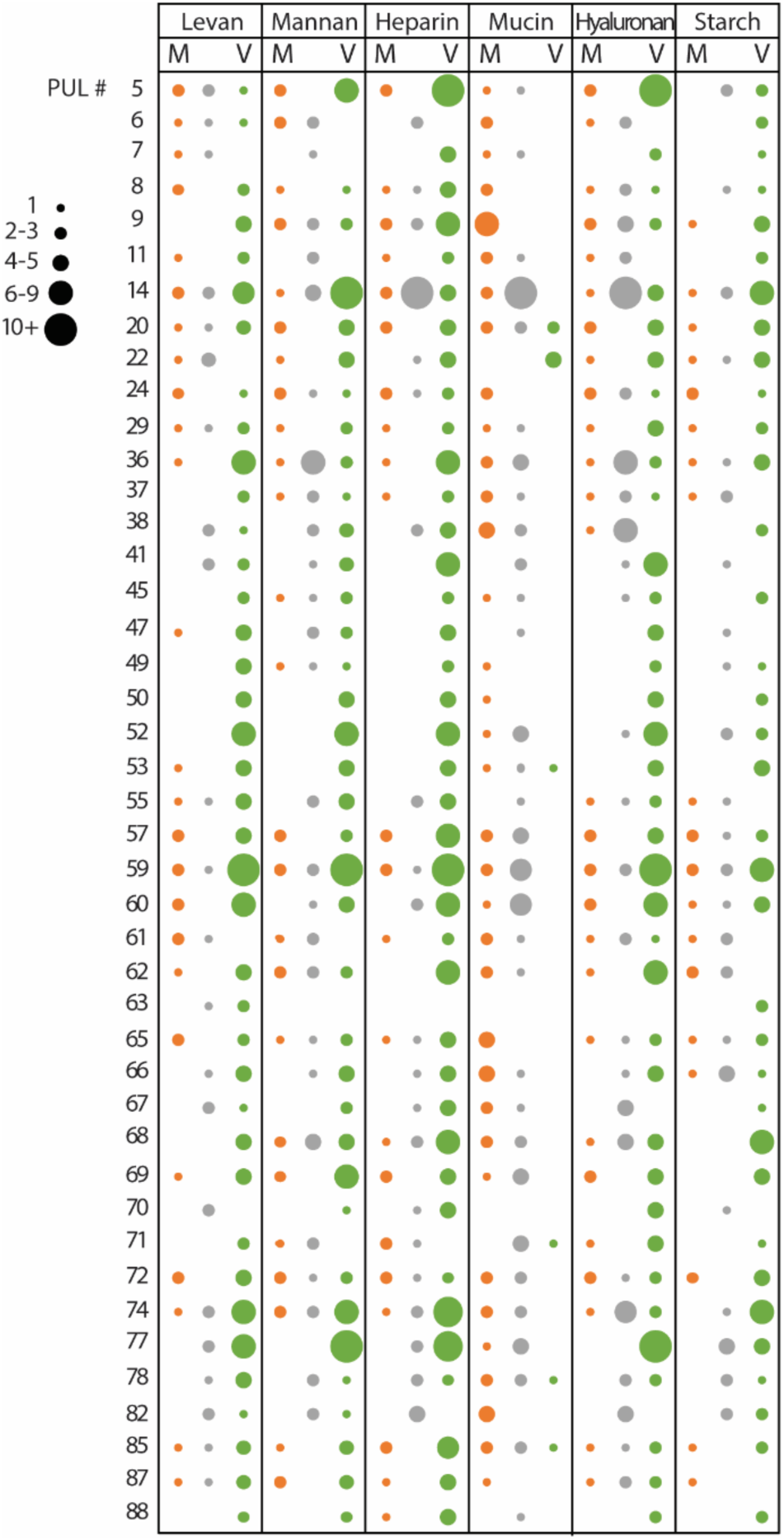
PUL-encoded proteins show OMV enrichment except for *Bt* grown in mucin. PUL-encoded proteins were identified and classified as OM-enriched (OMV/OM fold change <-1, M column, colored in orange), OMV-enriched (OMV/OM fold change >1, V column, colored in green) or unclassified (OMV/OM fold change between −1 and 1, colored in gray). Circle size represents the number of identified proteins for each PUL. Main representative PULs are shown (complete PUL dataset in figure S6).

Finally, we analyzed of the distribution of all predicted hydrolases between OMV and OM fractions, irrespective of their presence in PULs. Results revealed that about 60 to 80% of the total predicted hydrolases are enriched in OMV fractions in all the conditions, except for mucin, that exhibits only ∼20% of the total predicted hydrolases enriched in OMVs (fig. S8). Together, these results demonstrate that *Bt* can customize the OMVs protein cargo to optimize glycan utilization and suggest that vesicles are heavily involved in utilization of complex polysaccharides, but not in breakdown of mucin.

### OMVs liberate carbohydrate breakdown products supporting growth of other species

*Bacteroides* vesicles have been referred as “public goods” due to their ability to degrade extracellular polysaccharides whose breakdown products can be utilized by surrounding bacteria^23^. To functionally validate our proteomic findings, we analyzed the ability of OMVs to enhance the growth of other *Bacteroides* spp. in minimal media supplemented with different polysaccharides. We found that OMVs with cargo optimized for degradation of levan support the growth of poor levan utilizing bacteria, such as *Phocaeicola vulgatus* (*Pv,* reclassified from *Bacteroides vulgatus*^26^), *Bo* and *Bf*, in presence of levan. However, as predicted from our MS data, OMVs optimized for degradation of hyaluronan did not support growth of these bacteria in presence of levan (fig. 8). We also demonstrated the reciprocal, OMVs produced by *Bt* in hyaluronan support the growth of *Pv*, *Bo* and *Bf* in the same carbon source, but OMVs from *Bt* grown in levan do not. Similar results were obtained for other sugars (fig. 8). As suggested by our MS data, OMVs of *Bt* grown in mucin are not well-equipped to degrade mucin, and consistently, these did not improve growth of *Pv*, *Bo* or *Bf* in presence of mucin. Our results are consistent with the proposed roles of OMVs as public goods in the human gut^23^.

**Figure 8:**
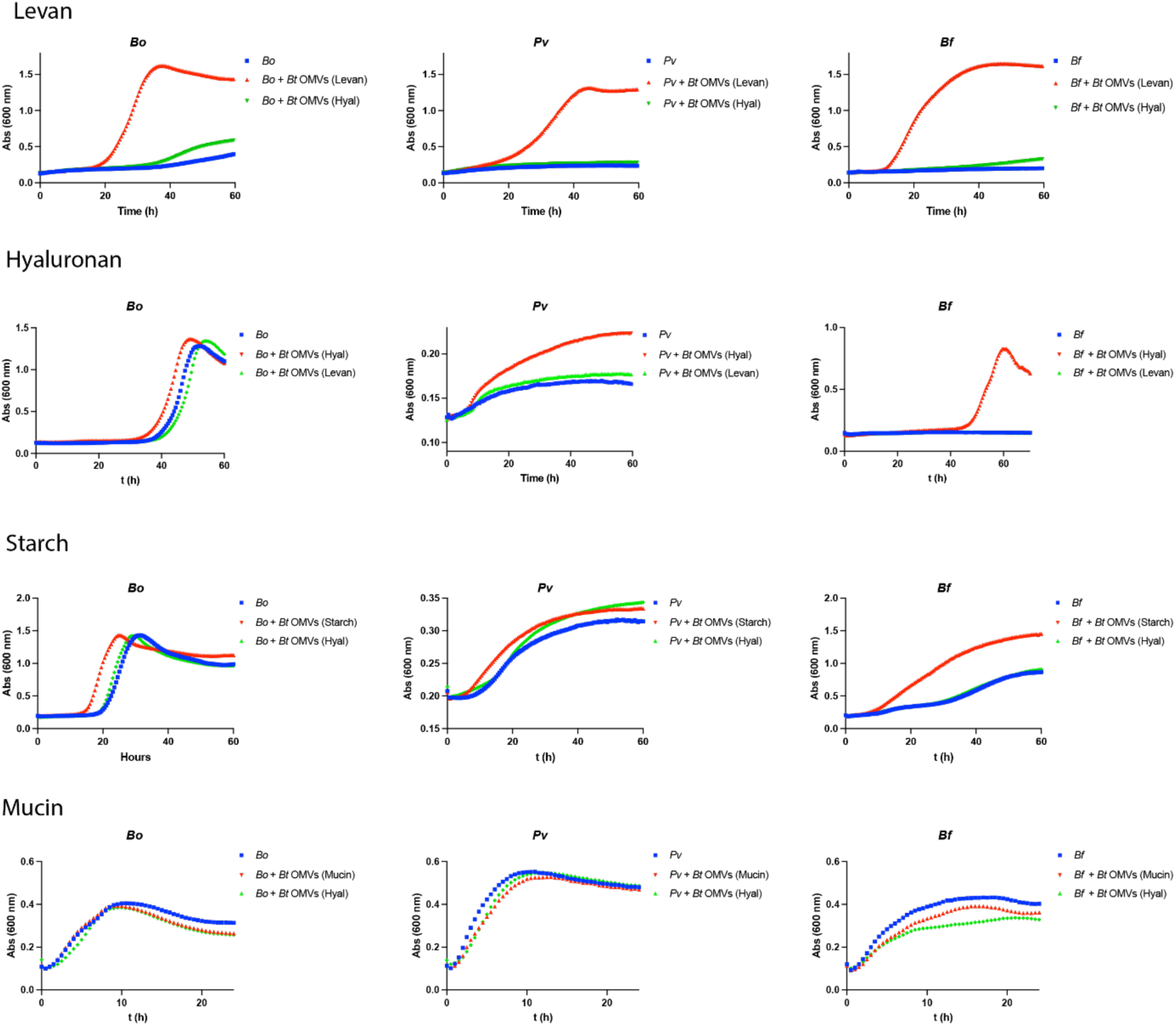
Tailored OMVs from *Bt* grown in a specific glycan enhance growth of *Bacteroides* spp. under the same culture condition. Growth curves of *Bo*, *Pv* and *Bf* in minimal media with levan, hyaluronan, starch, or mucin. Cultures were not supplemented (blue lines), supplemented with 1 μg/ml of *Bt* OMVs obtained after growth in the same glycan (red lines), or supplemented with 1 μg/ml of *Bt* OMVs obtained after growth in a different glycan (green lines).

## Discussion

In all three domains of life, cells secrete a large variety of membrane-bound extracellular vesicles that transport numerous molecules, and mediate intercellular communication in both physiological and pathological conditions^27^. In eukaryotes and archaea, EV biogenesis is known to rely on <u>E</u>ndosomal <u>S</u>orting <u>C</u>omplex <u>R</u>equired for <u>T</u>ransport (ESCRT) machinery^28–30^, which consists of a collection of highly conserved proteins that drive membrane budding and scission. However, despite the fast-growing literature related to vesicle production in bacteria, researchers have been unable to answer fundamental questions regarding how vesicle biogenesis occurs in bacteria, and how this process is regulated. In most Gram-negative bacterial species, OMVs and OM share very similar composition. In some cases, OMVs are reported to carry cytoplasmic components, such as ribosomal proteins, DNA, and RNA, that, according to the current knowledge of bacterial physiology, should not be packed into OMVs if vesicles are formed by blebbing of the OM. However, these elements can be associated with vesicles if they result from bacterial lysis. Indeed, some reports suggest that vesicles are formed by “explosive bacterial lysis”^31^, and most of the proposed mechanisms for OMVs biogenesis involve destabilization of the OM^32^. In consequence, the scientific community remains partially skeptical about the physiological relevance of OMVs. Based on our proteomic data, we have previously proposed the existence of a cargo selection process for *Bacteroides* vesicles^20, 21^. This unique characteristic of *Bacteroides*-derived OMVs prompted us to label bacteria and vesicles with different markers, allowing us to visualize, for the first time, vesicle biogenesis by live microscopy. Our microscopy analysis together with our exhaustive comparative proteomic analysis unequivocally demonstrate that OMVs are not the result of lysis, but a highly orchestrated process in *Bacteroides*. We also show that, based on a specific signal, the LES motif, OMV protein cargo is tailored to specifically degrade diet and host-derived polysaccharides. It remains to be demonstrated if similar dedicated OMV protein sorting mechanisms are present in other Gram-negative bacterial species.

Our observations led us to propose a model for vesicle production in *Bacteroides* (fig. 9). In the presence of a given polysaccharide, *Bacteroides* induces the expression of specific PULs required for binding, uptake and degradation of that glycan. Glycosyl hydrolases encoded in these PULs containing the LES signal are translocated across the OM, exposing them on the bacterial surface, where they cluster at defined foci in the OM. OMVs are subsequently generated by a still unknown machinery and subsequently released to the media. Production of OMVs enables the degradation of distal polysaccharides, and the resulting breakdown products can then be utilized by kin bacteria, other members of the microbiota, and even potential pathogens. This model does not exclude additional roles for OMVs in the context of interbacterial or host-bacteria interactions. In our microscopy experiments, we observed that OMV markers frequently localized at defined foci that were not detected in cells expressing OM markers (fig. 1). We propose that these foci represent regions of the OM where OMV proteins are recruited before the vesicles are released. It was also observed that bacterial cells do not always release the vesicles from these foci, which dynamically moved over time (fig. 3, supplementary video S1), in line with previous reported dynamic behavior for SusG^33^.

**Figure 9:**
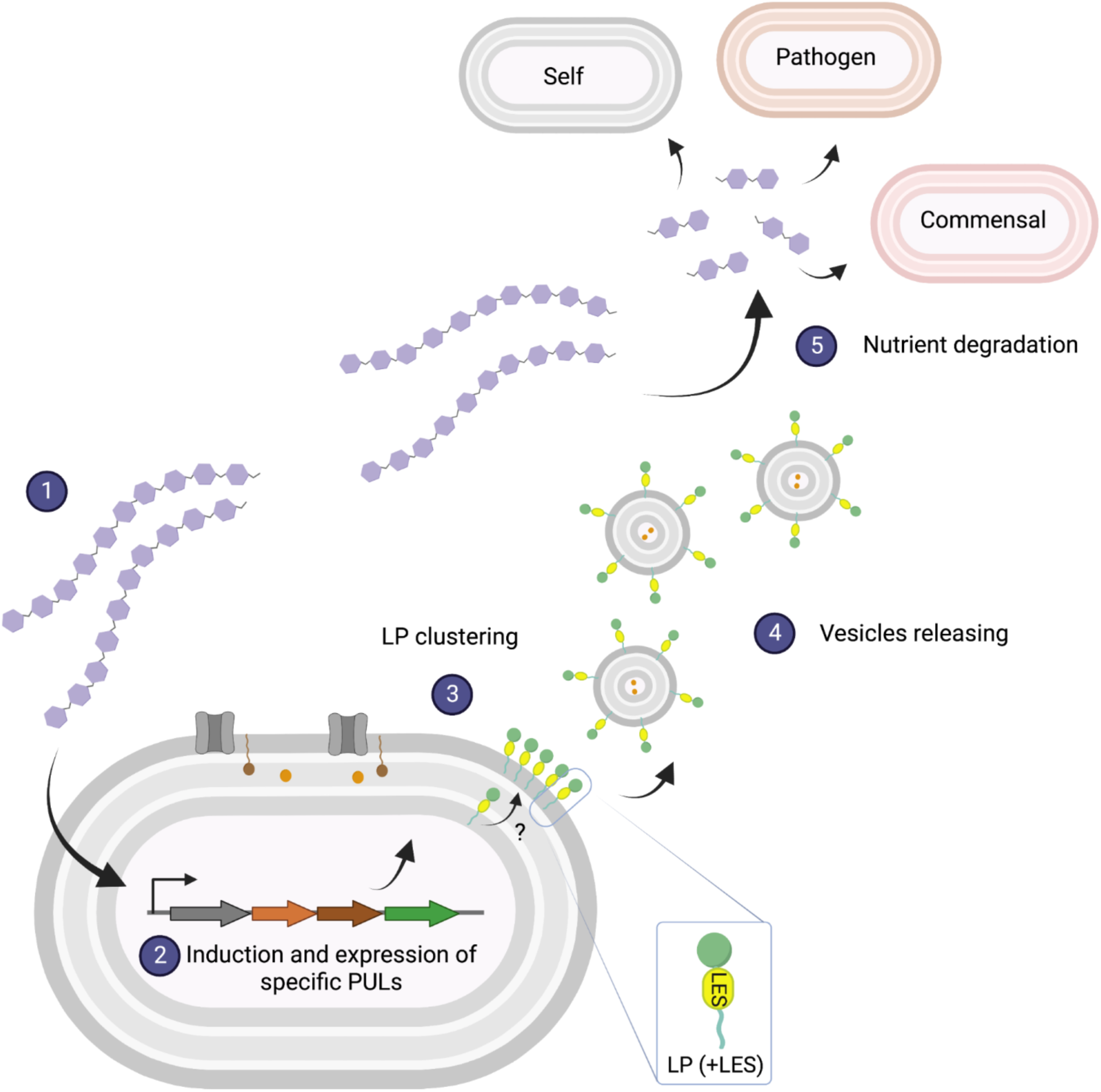
Model of OMV biogenesis. In the presence of a specific glycan (1), *Bacteroides* induces specific PULs (2). Glycosyl-hydrolases containing the LES signal are transported to the OM and exposed to the cell surface. OMV-targeted lipoproteins cluster at defined foci in the OM (3), where they are formed (4) by an unknown mechanism. OMVs diffuse and hydrolyze glycans that can be imported by the OMV-producing bacteria, as well as other commensals and pathogens (5). *Created with Biorender*.

Multiple comparative proteomic analyses revealed the existence of an underlying regulatory mechanism for OMV protein packing in *Bt*. We observed that, for each nutrient, a different subset of enzymes, several of them encoded in PULs, were packed into OMVs. In general, our proteomics data is in agreement with previous transcriptomic analyses that identified the PULs induced for the utilization of these glycans *in vitro* and *in vivo*^9^. However, we were able to demonstrate that an increase in the expression of PULs does not necessarily correlate with changes in the protein content of the OMVs. For each glycan analyzed in this work, most of the carbohydrate-induced enzymes containing the LES motif were targeted to OMVs, suggesting that OMVs play a role in their utilization. On the contrary, numerous proteins containing the LES motif were induced but retained in the OM in presence of mucin. We hypothesize that OMV biogenesis machinery is not induced in presence of mucin. Supporting these hypotheses, we observed significantly lower protein packed into OMVs when cultured in mucin as the sole carbon source (figures 4 and 5). Previous experiments performed in *Bt* in which mucin glycans were included together with other carbon sources show that degradation of mucin is not preferred in the presence of alternative substrates, even when *Bt* has been previously acclimated to growth in pure mucin glycans^34^. Altogether, these observations indicate that *Bacteroides* spp. have evolved to degrade mucus but as a last resource, and the fact that they pack less proteins into OMVs probably represents a strategy to avoid exacerbated degradation of the host intestinal mucus layer^35^.

Due to their size, OMVs diffuse beyond the bacterial surface, allowing the breakdown of nutrients that are out of reach for the producing bacteria. This is beneficial for the producing bacteria, but other microorganisms can also profit from OMV hydrolytic activities. Supporting this notion, *Bt* grown in starch displays a shorter lag phase when supplemented with physiological amounts of *Bt* OMVs produced in the same polysaccharide (fig. S9). In addition, OMVs can also enhance the growth of other species (fig. 7). An interesting example of cooperative feeding based on OMVs occurs between *Bo* and *Pv*. *Bo* secretes inulinases through OMVs, which supports the growth of *Pv*, unable to grow on inulin^23, 24^. Interestingly, it has been shown that the secreted and surface exposed inulinases of *Bo* are not required for its growth in inulin and, in fact, represent a fitness cost for the bacterium. However, this commensal relationship is reciprocal, as *Bo* benefits from *Pv* through an unknown mechanism. Also, pathogen-commensal interactions mediated via OMVs have been observed. The gastrointestinal human pathogen *Campylobacter jejuni* shows enhanced growth in mucin when co-cultured with *Pv* due to the secretion of fucosidases via OMVs^36^. Altogether, these examples reflect the co-evolution of commensal and pathogen species within the microbiota environment and highlight the diverse roles OMVs.

Our understanding of the physiological role of *Bacteroides* OMVs is still in its infancy. The molecular tools developed in this work, in particular the ability to distinguish between vesicles from cellular debris, will be essential to identify the molecular machinery responsible for OMV biogenesis. Additionally, this work lays the foundation for in-depth investigations into the biogenesis and function of *Bacteroides* vesicles within the human gut. Due to their important role in gut homeostasis, our research may lead, in the future, to the production of novel OMV-based therapies applicable to diseases involving gut dysbiosis.

## Materials and methods

### Bacterial strains and growth conditions

Strains, oligonucleotides, and plasmids are described in Table S1 in the supplemental material. *Bacteroides* strains were routinely grown in an anaerobic chamber (Coy Laboratories) using an atmosphere of 10% H_2_, 5% CO_2_, 85% N_2_. Bacteria were cultured in Brain heart infusion (BHI) medium supplemented with 5 μg/ml Hemin and 1 μg/ml vitamin K3. When applicable, antibiotics were used as follows: 100 μg/ml ampicillin, 200 μg/ml gentamicin and 25 μg/ml erythromycin. When required, *Bacteroides* was grown in minimal medium containing 100 mM KH_2_PO_4_ (pH 7.2), 15 mM NaCl, 8.5 mM (NH_4_)_2_SO_4_, 4 mM L-cysteine, 1.9 mM hematin/200 mM L-histidine (prepared together as a 1,000x solution), 100 mM MgCl_2_, 1.4 mM FeSO_4_.7H_2_O, 50 mM CaCl_2_, 1 μg/ml vitamin K3 and 5 ng/ml vitamin B12. Carbohydrates used to supplement MM include glucose (G7528, Sigma), levan (P-Levan, Neogen corp.), mannan (M7504, Sigma) starch (S2004, Sigma), hyaluronan (FH76335, Biosynth Carbosynth), heparin (YH09354, Biosynth Carbosynth) and mucin (M2378, Sigma). All carbon sources were added to MM in a final concentration of 0.5% (w/v).

### Growth assays

Bacterial strains were grown overnight in BHI, washed with PBS, and normalized by OD_600_ of 1. MM with different carbon sources were inoculated to a final OD_600_ of 0.05 and purified OMVs (1 μg/ml final concentration) were added when indicated. Growth curves were performed in sterile, round-bottom, polystyrene, 96-well plates in anaerobic and static conditions at 37 °C. OD_600_ values were recorded every 30 min with 10 s shaking before measurements with SmartReader MR9600-T microplate reader. All experiments were performed on 3 independent days with at least 3 wells per strain per condition.

### Creation of protein markers fused to GFP and mCherry

OMVs (BT_3968, Bacova_04502, and BF_1581) and OM (BT_0418 and BT_2844) enriched proteins selected as markers were fused to sfGFP and/or mCherry. For this, the genes codifying for the proteins of interest were amplified by PCR using primers listed in table S1. The purified products were cloned into pWW plasmid series upstream of sfGFP/mCherry coding sequences^37^ using NEBuilder® HiFi DNA Assembly (NEB). To perform co-expression experiments, OMV and OM fluorescent chimeric markers were cloned into the same integrative plasmid. Constructs were conjugated into *B. thetaiotaomicron* using previously transformed *Escherichia coli* S17-1 λ*pir* as a donor. Strains were plated and *Bacteroides* strains harboring the integrated plasmid were selected in BHI agar plates supplemented with 200 μg/ml gentamicin and 25 μg/ml erythromycin.

### Vesicles preparations

OMVs were purified by ultracentrifugation of filtered spent media. Briefly, 50 ml of of *B. thetaiotaomicron* cultures from early stationary phase were centrifuged at 6,500 rpm at 4 °C for 10 minutes. Supernatants were filtered using a 0.22-μm-pore membrane (Millipore) to remove any residual cells. The filtrate was subjected to ultracentrifugation at 200,000 x*g* for 2 h (Optima L-100 XP ultracentrifuge; Beckman Coulter). Supernatants were discarded, and the pellet resuspended in PBS. Purified OMV preparations were lyophilized for MS analysis.

### Subcellular fractionation

Total membrane preparations were performed by cell lysis and ultracentrifugation. Cultures from early stationary phase were harvested by centrifugation at 6,500 rpm at 4°C for 10 minutes. The pellets were gently resuspended in a mixture of 50 mM Tris-HCl (pH 8.0), 150 mM NaCl, and 50 mM MgCl_2_ containing complete EDTA-free protease inhibitor mixture (Roche Applied Science) followed by cell disruption. Centrifugation at 6,500 rpm at 4 °C for 5 minutes was performed to remove unbroken cells. Total membranes were collected by ultracentrifugation at 200,000 x*g* for 1 h at 4 °C, and the soluble fraction was collected from the supernatant. OM and IM were separated by differential extraction with the same buffer supplemented with 1% (v/v) *N*-lauroyl sarcosine and incubated 1 h at room temperature with gentle agitation. The OM fractions were recovered by centrifugation at 200,000 x*g* for 1 h at 4 °C in the pellet fraction, whereas the IM fraction was obtained from the supernatant. OM fractions were lyophilized for MS analysis.

### SDS-page and Western blot analyses

Membrane and vesicles fractions were analyzed by standard 10% Tris-glycine SDS-PAGE. Briefly, 10 μg of each fraction was loaded onto an SDS-PAGE gel and stained with Coomasie blue for total protein analysis, or transferred onto a nitrocellulose membrane for Western blot analysis. Membranes were blocked using Tris-buffered saline (TBS)-based Odyssey blocking solution (LI-COR). Primary antibodies used in this study were rabbit polyclonal anti-His (ThermoFisher), and rabbit polyclonal anti-mCherry (ThermoFisher). Secondary antibodies used were IRDye anti-rabbit 780 (LI-COR). Imaging was performed using an Odyssey CLx scanner (LI-COR).

### Widefield fluorescence microscopy

Bacteria were swabbed from BHI agar plates into MM supplemented with 0.5% (w/v) of glucose and cultured for 20 hs in anaerobic chamber at 37 °C. For GFP and mCherry fluorophores maturation^38–40^, cultures were removed from the anaerobic chamber and incubated at 37 °C for 4 h in aerobic conditions. Fifty microliters of bacteria were diluted into 200 μl PBS and 3 μl of the dilution was dotted onto 1% agarose pads in PBS. Excess of liquid was air-dried and 18mm coverslips were added. Images were acquired using a Zeiss Axio Imager M2 upright fluorescence microscope equipped with Plan apo 100x/1.40 N.A. Oil Ph3 M27 objective lens. Images were adjusted and cropped using Fiji^41^.

For time lapse fluorescence microscopy, cell growth and fluorophore maturation were performed as previously described. Fifty microliters of bacteria were diluted into 200 μl of pre-warmed MM supplemented with glucose and 3 μl of the dilution was dotted onto 35 mm glass bottom culture dishes (MatTek). Drop was covered with prewarmed 1% agarose pads in MM supplemented with glucose. Images were acquired using a Zeiss Cell Observer Z1 inverted microscope equipped with a temperature-controlled incubation chamber at 37°C. Fluorescence images were acquired every 3.5 min with illumination from a Colibri 7 LED light source (Zeiss) and ORCA-ER digital camera (Hammamatsu Photonics, Japan). A Plan-Apochromat 63x/1.4 N.A. Phase 3 objective and ZEN blue 2.5 software were used for image acquisition.

### Protein sample preparation for Mass Spectrometry analyses

Lyophilized protein preparations were solubilized in 100 μl of 5% SDS by boiling them for 10 min at 95 °C. The protein content was assessed by the bicinchoninic acid (BCA) protein assay (Thermo Fisher Scientific) according to manufacturer’s instructions. One hundred microgram of each sample were reduced with 10mM DTT for 10 mins at 95°C and alkylated with and alkylated with 40mM IAA in the dark for 1 hour. Reduced and alkylated samples were cleaned up using Micro S-traps (https://protifi.com/pages/s-trap) according to the manufacturer’s instructions. Samples were digested overnight with 3 μg of trypsin/Lys-C (1:33 protease/protein ratio) and then collected. Samples were dried down and further cleaned up using C18 Stage tips^42, 43^ to ensure the removal of any particulate matter.

### LC-MS

Prepared purified peptides from each sample were re-suspended in Buffer A* (2% acetonitrile, 0.01% trifluoroacetic acid) and separated using a two-column chromatography setup composed of a PepMap100 C_18_ 20-mm by 75-μm trap and a PepMap C_18_ 500-mm by 75-μm analytical column (Thermo Fisher Scientific). Samples were concentrated onto the trap column at 5 μl/min for 5 min with Buffer A (0.1% formic acid, 2% DMSO) and then infused into an Orbitrap Q-Exactive plus Mass Spectrometer (Thermo Fisher Scientific) or a Orbitrap Fusion Lumos equipped with a FAIMS Pro interface at 300 nl/minute via the analytical column using a Dionex Ultimate 3000 UPLC (Thermo Fisher Scientific). Ninety-five-minute analytical runs were undertaken by altering the buffer composition from 2% Buffer B (0.1% formic acid, 77.9% acetonitrile, 2% DMSO) to 22% B over 65 min, then from 22% B to 40% B over 10 min, then from 40% B to 80% B over 5 min. The composition was held at 80% B for 5 min, and then dropped to 2% B over 2 min before being held at 2% B for another 8 min. The Q-Exactive plus Mass Spectrometer was operated in a data-dependent mode automatically switching between the acquisition of a single Orbitrap MS scan (375-1400 m/z, maximal injection time of 50 ms, an Automated Gain Control (AGC) set to a maximum of 3 x 10^6^ ions and a resolution of 70k) and up to 15 Orbitrap MS/MS HCD scans of precursors (Stepped NCE of 26%, 28% and 32%, a maximal injection time of 110 ms, an AGC set to a maximum of 2 x 10^5^ ions and a resolution of 35k). The Fusion Lumos Mass Spectrometer was operated in a stepped FAIMS data-dependent mode at two different FAIMS CVs −45 and −65. For each FAIMS CV a single Orbitrap MS scan (300-1600 m/z, maximal injection time of 50 ms, an AGC of maximum of 4 x10^5^ ions and a resolution of 60k) was acquired every 1.5 seconds followed by Orbitrap MS/MS HCD scans of precursors (NCE 35%, maximal injection time of 100 ms, an AGC set to a maximum of 1.25 x10^5^ ions and a resolution of 30k).

### MS data analysis

Identification and LFQ analysis were accomplished using Max-Quant (v2.0.2.0) ^44^ using *Bacteroides thetaiotaomicron* VPI-5482 proteome (Uniprot: UP000001414) allowing for oxidation on Methionine. Prior to MaxQuant analysis dataset acquired on the Fusion Lumos were separated into individual FAIMS fractions using the FAIMS MzXML Generator^45^. The LFQ and “Match Between Run” options were enabled to allow comparison between samples. The resulting data files were processed using Perseus (v1.4.0.6)^46^ to filter proteins not observed in at least four biological replicates of a single group. ANOVA and Pearson correlation analyses were performed to compare groups. Predicted localization and topology analysis for proteins identified by MS were performed using UniProt^47^, PSORT^48^ and SignalP^49^.

## Data availability

The mass spectrometry proteomics data have been deposited in the Proteome Xchange Consortium via the PRIDE partner repository with the data set identifier PXD036181 (reviewer login details: **Username:****Password:** zIcssrKS); PXD036275 (reviewer login details: **Username:****Password:** ezaiiNkH) and PXD036272 (reviewer login details: **Username:****Password:** G4q7jZDD)^50^.

## Acknowledgments

We thank all members of the Feldman lab for critically reading the manuscript. We also thank Wandy Beatty for assistance with fluorescence microscopy experiments.

## Authors contribution

MFF, MGS and EJP designed and wrote the manuscript. MGS and EJP performed experiments on *Bacteroides*. NES performed MS experiments and data analysis.

---

**Figure S1.**
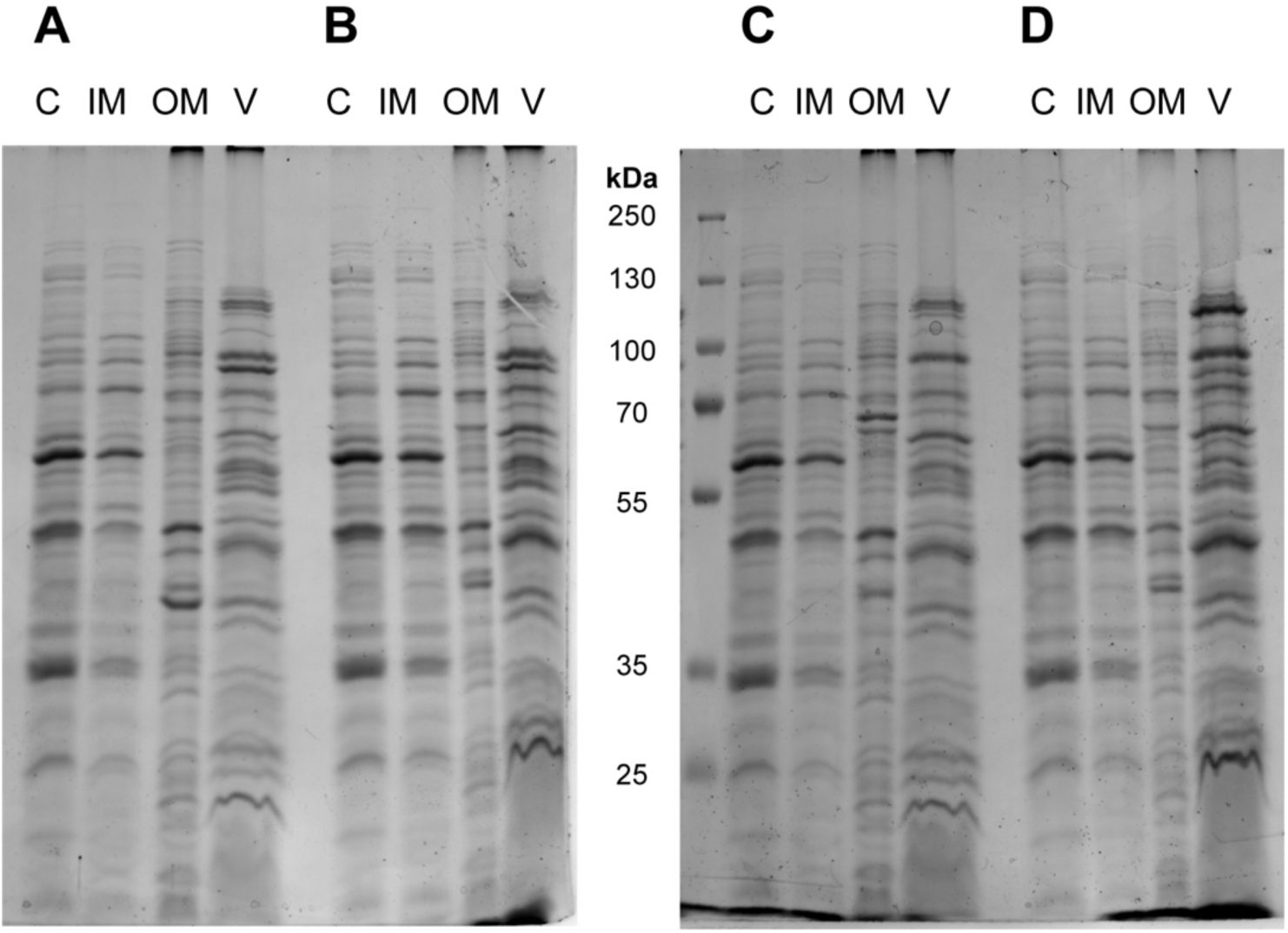
Coomassie blue staining after SDS-PAGE of subcellular fractionations (10 μg) of *Bt* expressing **A.** Inulinase-GFP, **B.** BF_1581-GFP, **C.** OmpF-mCherry, and **D.** BT_2844-mCherry. References: S, soluble fraction; IM, inner membrane fraction; OM, outer membrane fraction; V, OMV fraction.

**Figure S2.**
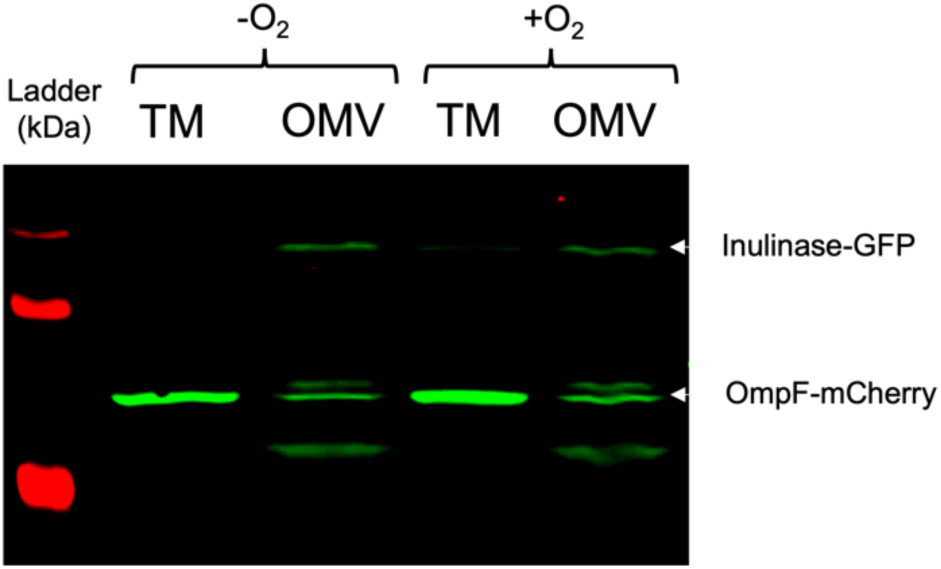
Western blots after SDS-PAGE of total membranes (TM) and OMVs from *Bt* co-expressing Inulinase-GFP and OmpF-mCherry. Bacteria were swabbed from BHI agar plates into MM supplemented with 0.5% (w/v) of glucose and cultured for 20 hs in anaerobic chamber at 37 °C. Cultures were then maintained in anaerobiosis (-O_2_) or exposed for 4 hours to atmospheric conditions (+O_2_) prior to fractionation. Anti-His and anti-mCherry antibodies were employed to identify GFP and mCherry chimeric markers, respectively.

**Figure S3.**
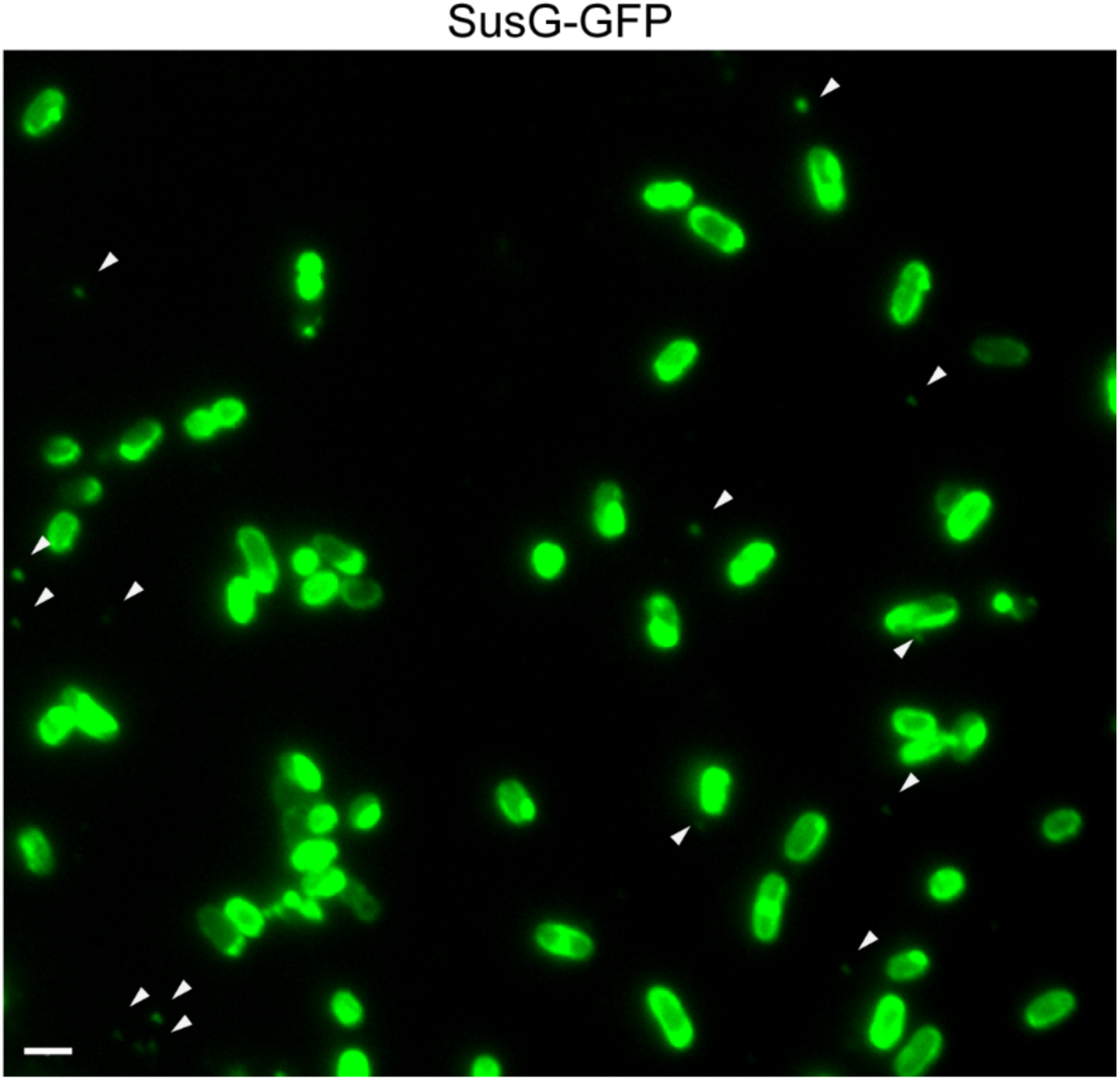
Representative widefield fluorescence microscopy images of OMV chimeric marker SusG-GFP showing OMVs and localization at defined foci on the bacterial surface. Scale bar: 2 μm.

**Figure S4.**
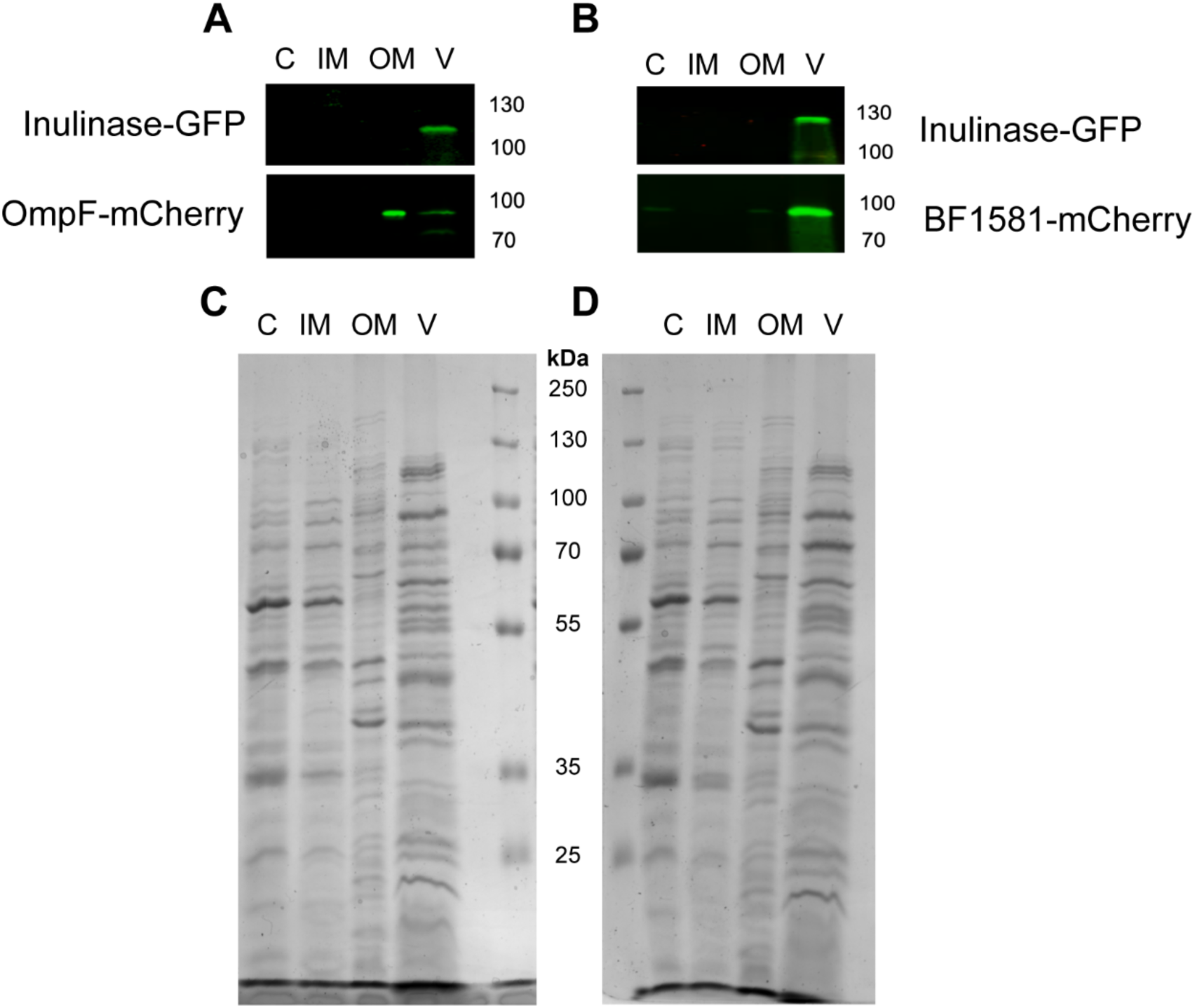
Western blots and Coomasie blue staining after SDS-PAGE of subcellular fractions of *Bt* co-expressing Inulinase-GFP and OmpF-mCherry (A and C, respectively), or Inulinase-GFP and BF_1581-GFP (B and D, respectively). References: soluble fraction (S), inner membrane (IM), outer membrane (OM) and OMVs (V). Anti-His and anti-mCherry antibodies were employed to identify GFP and mCherry chimeric markers, respectively.

**Figure S5.**
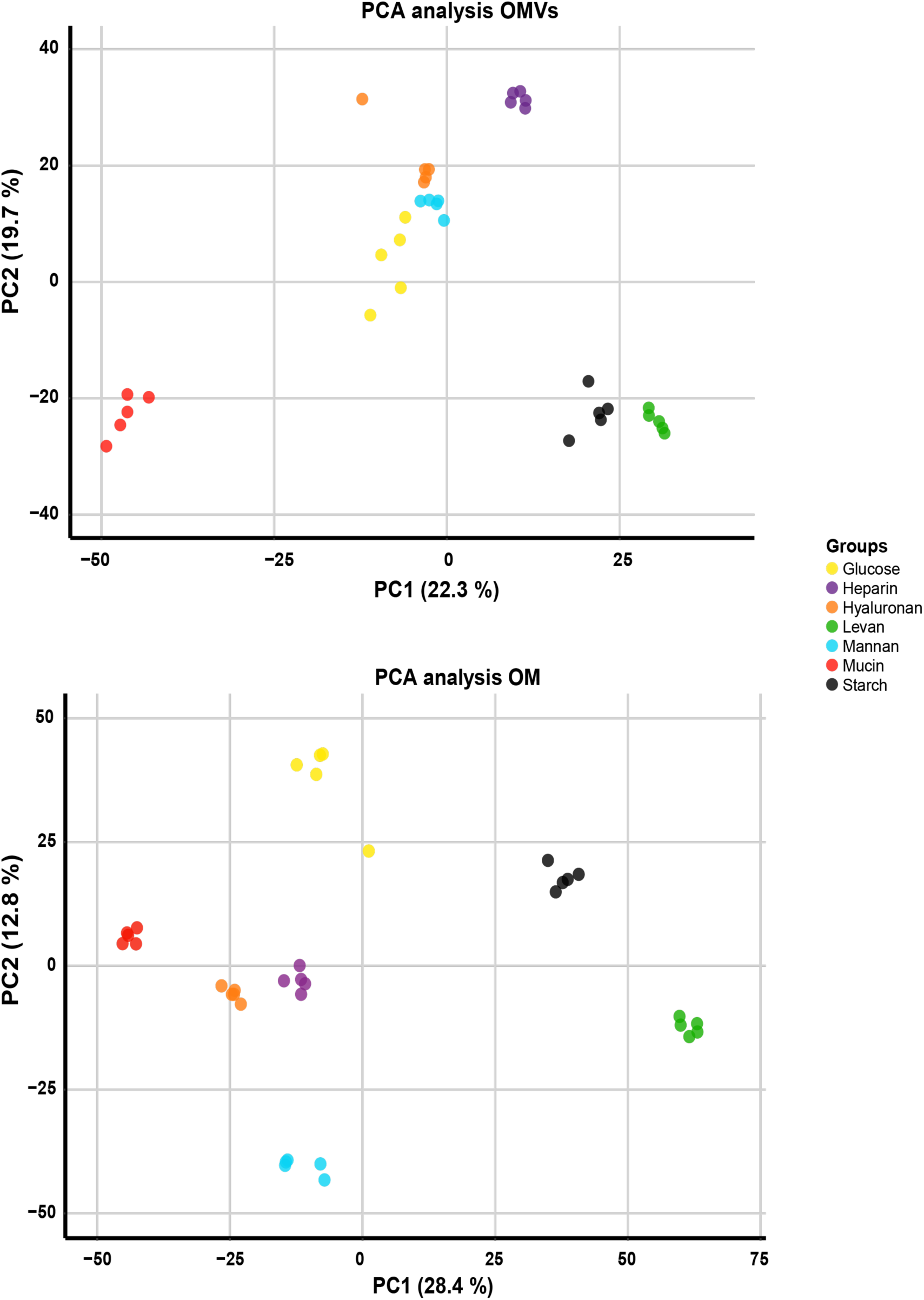
Principal component analysis (PCA) of OMV (upper panel) and OM proteomes (lower panel) from *Bt* grown in minimal media supplemented with the indicated carbon sources. Five biological replicates were performed for each condition.

**Figure S6.**
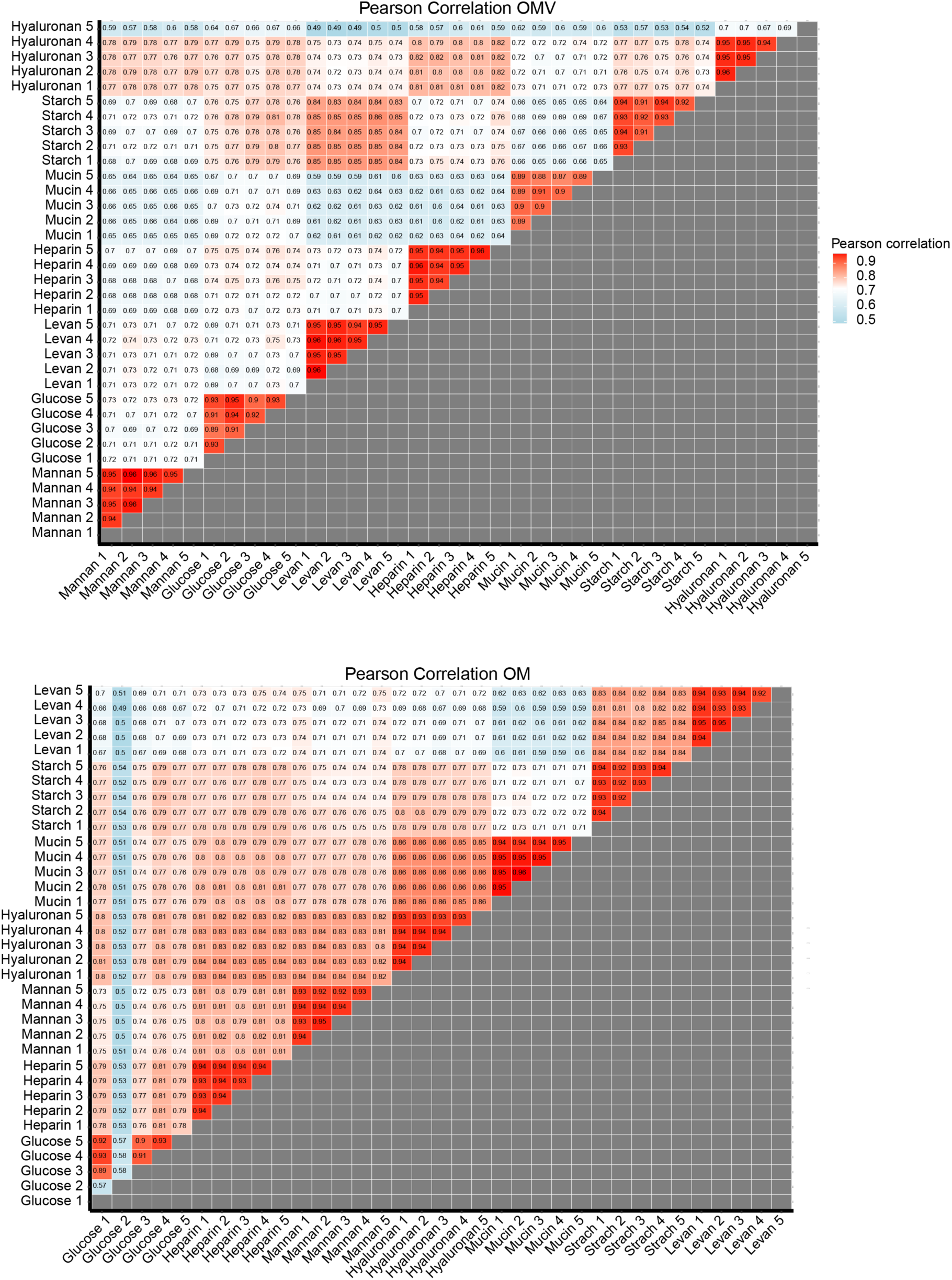
Pearson Correlation between OMV (upper panel) and OM samples (lower panel) from *Bt* grown in minimal media supplemented with the indicated carbon sources. Five biological replicates were performed for each condition.

**Figure S7.**
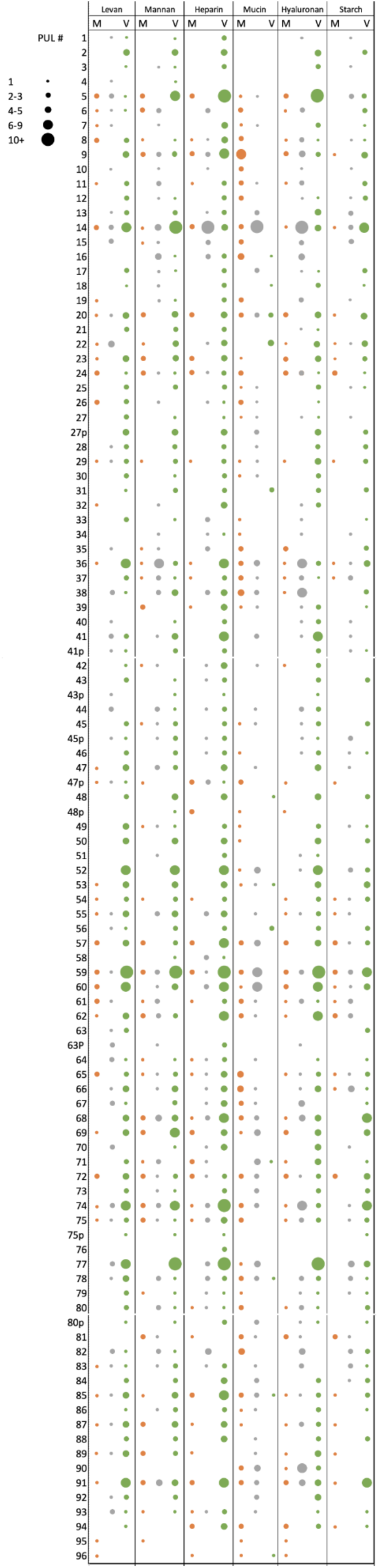
PUL-encoded proteins were identified and classified as OM-enriched (OMV/OM fold change <-1, M column, colored in orange), OMV-enriched (OMV/OM fold change >1, V column, colored in green) or unclassified (OMV/OM fold change between −1 and 1, colored in gray). Circle size represents the number of identified proteins for each PUL.

**Figure S8.**
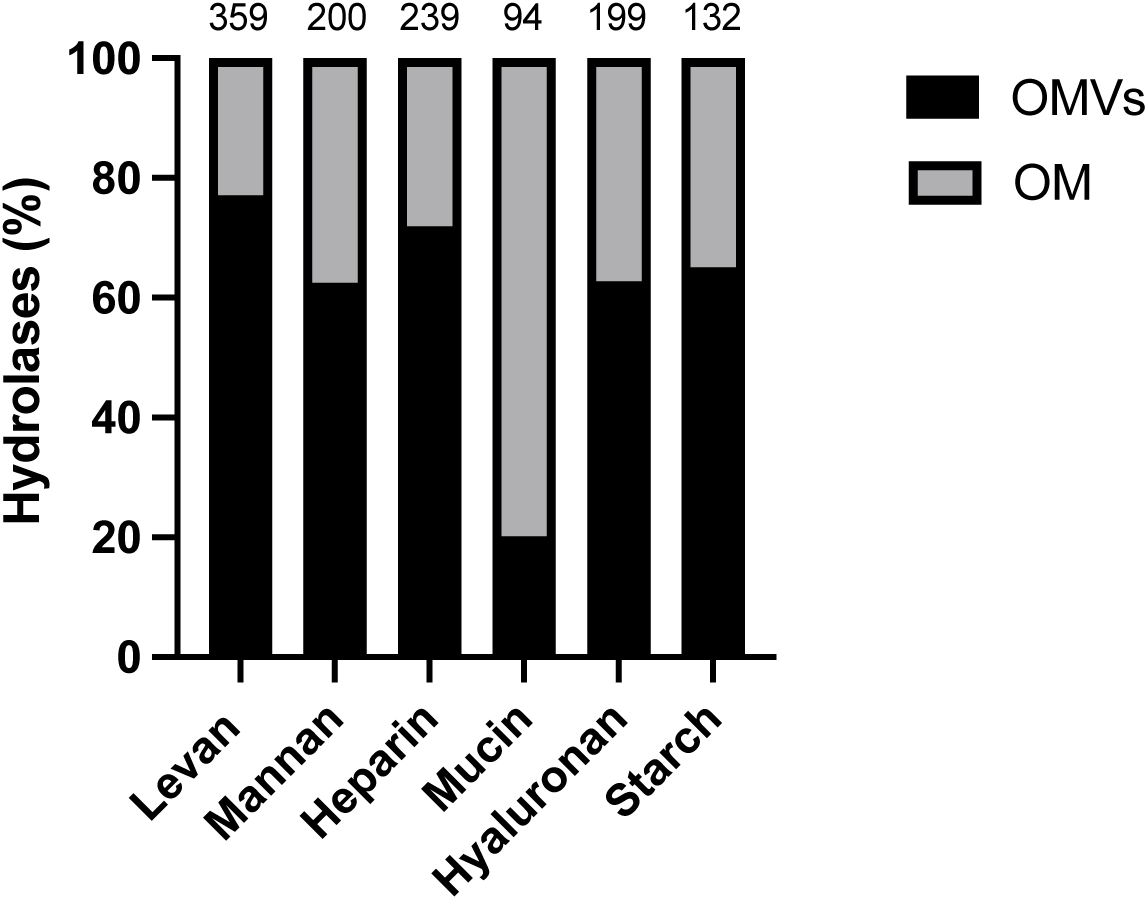
Hydrolases partitioning between OM and OMV fractions. All predicted hydrolases identified for each condition were classified as OM-enriched (OMV/OM fold change <-1) or OMV-enriched (OMV/OM fold change >1). Numbers at the top of bars indicate the total number of hydrolases identified in each condition.

**Figure S9.**
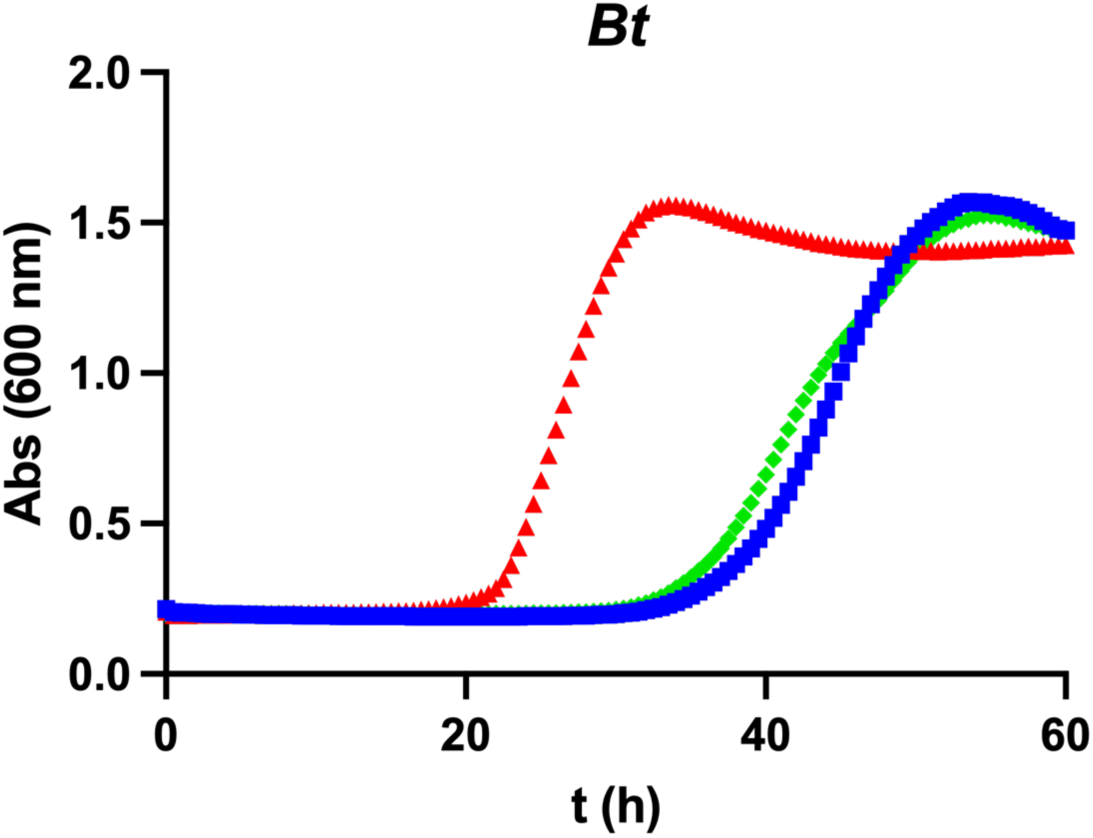
Growth curves of *Bt* in minimal media with starch. Cultures were not supplemented (blue lines), supplemented with 1 μg/ml of *Bt* OMVs obtained after growth in the same glycan (red lines), or supplemented with 1 μg/ml of *Bt* OMVs obtained after growth in a different glycan (hyaluronan, green lines).

**Table S1.** Strains, plasmids and oligonucleotides used in this study.

**Table S2.** Comparative proteomic analysis between OMVs and OM fractions from Bt grown in MM supplemented with levan.

**Table S3.** Comparative proteomic analysis between OMVs and OM fractions from *Bt* grown in MM supplemented with mannan.

**Table S4.** Comparative proteomic analysis between OMVs and OM fractions from Bt grown in MM supplemented with heparin.

**Table S5.** Comparative proteomic analysis between OMVs and OM fractions from *Bt* grown in MM supplemented with mucin.

**Table S6.** Comparative proteomic analysis between OMVs and OM fractions from Bt grown in MM supplemented with hyaluronan.

**Table S7.** Comparative proteomic analysis between OMVs and OM fractions from *Bt* grown in MM supplemented with starch.

**Table S8.** Comparative proteomic analysis between OMVs and OM fractions from Bt grown in MM supplemented with glucose.

**Table S9.** Comparative proteomic analysis between OMVs from all the growth conditions showing most enriched proteins in each carbon source.

**Table S10.** Comparative proteomic analysis between OMs from all the growth conditions showing most enriched proteins in each carbon source.

**Supplementary Movie S1.** Live OMV formation.

